# Near-anoxia induces immobilisation and sustains viability of sperm in ant queens: Novel insights into mechanisms of prolonged sperm storage

**DOI:** 10.1101/2022.09.08.507031

**Authors:** Ayako Gotoh, Mika Takeshima, Ken-ichi Mizutani

## Abstract

Insect females store sperm in a spermatheca after copulation for some duration to use it later for fertilisation. At the beginning of their adult lives, ant queens can preserve numerous viable sperm cells from copulation for over ten years. However, the key factors influencing long-term sperm storage have not been identified. Here we show that the spermathecal environment is near anoxic, which induces sperm immobilisation. Furthermore, mitochondrial respiratory inhibitors suppress sperm motility, suggesting that sperm immobilisation may be caused by a shortage of ATP generated from only glycolysis under near anoxic condition. Sperm immobilisation is not induced by acidification via glycolytic metabolism because the spermathecal fluid is not acidic. Finally, we show that artificial anoxic conditions rather than aerobic conditions sustain viable sperm cells. Therefore, near-anoxia is a key factor influencing long-term sperm storage in ant queens. The viability of sperm cells under artificial anoxia, however, is lower than of those dissected immediately from queens. Moreover, the immotile sperm cells under more than 4 h of anoxia do not begin swimming after aerobic exposure, unlike those that were in anoxic conditions for less than 2 h. This indicates that factors other than anoxia are also necessary for long-term sperm preservation.

## 1. Introduction

Various unique reproductive strategies for reproductive success are known in both sexes of sexual organisms. Among them, many species of various taxa have evolved the strategy of sperm storage between copulation and fertilisation inside the female reproductive tract [1]. The duration of sperm storage ranges from temporary to over a decade among various species. Ant queens copulate only at the beginning of their adult lives and use the sperm for fertilisation until they die; therefore, they can maintain viable sperm cells for the same duration as their life span. Because queens can live for more than ten years in many ant species [2], their capacity for sperm maintenance is pronounced. However, the physiological mechanisms by which ant queens maintain viable sperm cells are still unclear.

Ant queens maintain sperm cells inside their sperm storage-sac (spermathecal reservoir) [3]. It consists of mitochondria-rich columnar epithelia and flattened epithelia with few organelles [3, 4]. The flattened epithelium is considered to not affect the sperm storage function because of its ultrastructure. In contrast, the columnar epithelium is considered to perform a transporting function between inside and outside of the spermathecal reservoir. It has been reported that sperm cells stored in the spermathecal reservoir are immobilised [5]. Furthermore, the immobilisation has also been observed during sperm transfer from the bursa copulatrix to the spermathecal reservoir via the spermathecal duct soon after copulation [5]. Sperm immobilisation is considered to have significant advantages for long-term sperm storage in avoiding the risk of sperm damage and inhibiting the production of reactive oxygen species if they use oxygen to produce adenosine triphosphate (ATP). Therefore, understanding the mechanism of sperm immobilisation could elucidate the mechanism of prolonged sperm storage in ant queens.

Many studies have reported on factors that induce sperm motility after male ejaculation in many animals, e.g., serine proteases in silkworms [6], and change in extracellular osmolality inducing a change in intracellular ion concentration in many teleost species [7]. However, studies on the regulatory mechanisms that regulate the motility of sperm stored in females are limited [8].

In this study, we show that near anoxia in the spermathecal reservoir is a key factor influencing sperm immobilisation, and artificial anoxia rather than aerobic conditions can sustain viable sperm cells.

## 2 Material and Methods

### (a) Ants

We collected *Crematogaster osakensis* and *Lasius hayashi* queens from the ground after nuptial flights in Kagawa and Hyogo prefectures in western Japan. The queens were housed in plastic cases with moistened plaster bases at 25 °C. *L. hayashi* queens were used only for quantifying oxygen concentration, and *C. osakensis* queens at three to ten months after mating were used for sperm motility and viability experiments. We used queens of the same age and from the same collection site in the comparative experiments.

### (b) Measurement of oxygen concentration

We measured oxygen concentrations in the spermathecal reservoir, and thoracic and abdominal haemolymph of *C. osakensis* queens at one year and *L. hayashi* queens at seven months after the beginning of sperm storage (i.e., nuptial flight) using a calibrated TX3 trace oxygen microoptode (PreSens). The spermatheca was dissected from the queen, and the oxygen probe was inserted into the spermathecal reservoir. The oxygen probe was also inserted into the thorax and abdomen through a small hole created in the intersegment using forceps. To measure oxygen concentrations in *C. osakensis* queens soon after copulation, we dissected queens from 9 to 10 p.m. on the night of the nuptial flight [5].

### (c) Observation of sperm motility under artificial hypoxic conditions using an O_2_-absorbing agent

*C. osakensis* sperm cells extracted from the spermathecal reservoir are known to begin swimming in phosphate-buffered saline (PBS) [5]. To create an artificial hypoxic test solution, an 8- well chamber slide containing 800 μL Dulbecco’s PBS without calcium and magnesium (Nacalai Tesque) and an AnaeroPack^®^ kenki pouch (Mitsui Gas Co.), which absorbs O_2_ and generates CO_2_, were kept in a sealed plastic bag for 24 h. After 24 h, the oxygen content in the PBS was 0.4–0.5% (n = 3), and the pH had not changed significantly. The sperm cells from the spermatheca of *C. osakensis* queen were exposed to the hypoxic PBS. Furthermore, PBS in an 8-well chamber slide placed in atmospheric conditions for 24 h was used as control. The movie of the sperm motility was captured using an inverted microscope (Olympus CKX53) and 3CCD digital camera (Olympus DP72). To avoid atmospheric oxygen dissolution in the artificial hypoxic PBS, sperm ejection from the spermatheca and motility observation was performed as quickly as possible (within 2 min). However, it is likely that sperm cells exposure to atmospheric oxygen varied slightly because the time it took to introduce the sperm cells to the solution differed between samples, and oxygen immediately contaminates the PBS on exposure to the atmosphere. Moreover, because the effect of CO_2_ (generated from AnaeroPack kenki pouch) on sperm immobilisation could not be eliminated, we used a test solution containing sodium sulphite as an alternative method to create an artificial anoxic environment (section 2(d)).

### (d) Effect of oxygen scavenger, inhibitors of glycolysis and respiration, and pH on sperm motility

To investigate the effect of anoxia on sperm immobilisation, we prepared 39.44 mM sodium sulphite (Fujifilm Wako Pure Chemical Corporation), as an oxygen scavenger (anoxic condition), in PBS. We also prepared an oxygen-saturated sodium sulphite solution in PBS (aerobic condition) by leaving it for a day. The oxygen concentrations in the solutions were verified using a TX3 trace oxygen microoptode. We also prepared 10 μM antimycin (Enzo Life Sciences), oligomycin (Abcam), and carbonyl cyanide 4-(trifluoromethoxy) phenylhydrazone (FCCP) (Cayman chemical) in PBS as inhibitors of the mitochondrial respiratory pathway. In addition, as glycolytic pathway inhibitors, we prepared 50 μM heptelidic acid (Abcam) and 1 mM iodoacetic acid (Fujifilm Wako Pure Chemical Corporation) in PBS. To investigate effect of pH on sperm motility, 150 mM NaCl with pH 7.1 and 8.6 by adding NaOH and HCl, respectively were also prepared. To prevent change in pH, the NaCl solution was used within 5 min of pH adjustment for each experiment.

A 0.5-mm deep silicone rubber spacer was placed on a slide to keep space between the slide and coverslip. Sperm cells were exposed to 20 μL of test solution on the slide and covered with a coverslip. The slide was observed and photomicrographed.

Sperm motility levels were scored based on the frequency of motile sperm cells as follows: 0, no motile sperm cells; 1, some sperm cells exhibiting motility; 2, approximately 40% sperm cells exhibiting motility; 3, 80–90% motility; and 4, maximum sperm cells motility. The scoring was performed by a trained observer within 2 min of the dissection.

### (e) Determining pH of spermathecal fluid and sperm

The pH of spermathecal fluid from spermatheca of *C. osakensis* queen was measured using pH paper (pH range 5.5–9.0; Macherey-Nagel). The pH of the haemolymph from a cut on the trochanter of the hind leg was also measured as a control.

To assess the influence of extracellular fluid pH on the sperms’ intracellular pH, the sperm cells were stained with BCECF-AM (Dojindo) intracellular pH indicator diluted 1000-fold with pH calibrated buffers, 4.5, 5.5, 6.5, and 7.5 (Intracellular pH Calibration Buffer Kit, Thermo Fisher Scientific), and with the staining buffers containing 10 μM valinomycin and 10 μM nigericin to equilibrate the pH inside and outside of cells, for 10 min. After the sperm samples were centrifuged at 5000 rpm for 5 min and the supernatants were removed, a further 20 μL of each buffer was added. From this suspension, a 3 μL aliquot was deposited on a slide, covered with a coverslip, and observed under a fluorescence microscope (Olympus BX53 combined with U-FGW filters) and 3CCD digital camera (Olympus DP74) were used to capture photomicrographs.

### (f) Determining of ATP levels in sperm

To determine intracellular ATP, sperm cells were introduced to 40 μL of 39.44 mM sodium sulphite in PBS (anoxic condition) and PBS (aerobic condition) for 1 min. Thereafter, the sperm samples were centrifuged at 5000 rpm for 5 min, after which the supernatants were removed, and CellTiter-Glo^®^ Luminescent Cell Viability Assay buffer (Promega) was added. The sperm suspension was then divided into two samples: one for determining ATP using a CellTiter-Glo Luminescent Cell Viability Assay kit (Promega) and the other for assessing DNA content (to calibrate the ATP content by the number of sperm cells) using CellTox^™^ Green Cytotoxicity Assay kit (Promega). After the samples were prepared according to the manufacturer’s protocol, their luminescence and fluorescence were detected using the GloMax^®^ system (Promega).

### (g) Assessing sperm mitochondrial activity

To assess the membrane potential of sperm mitochondria, sperm cells were stained using 40 μL of an MT-1 MitoMP Detection Kit (Dojindo), diluted 1000-fold with 39.44 mM sodium sulphite in PBS (anoxic condition), PBS (aerobic condition) and 10 μM FCCP in PBS (negative control) for 15 min. After the sperm samples were centrifuged at 5000 rpm for 5 min and the supernatants were removed, a further 20 μL of each solution was added. Further, 3 μL of the sperm suspension was placed on a slide, covered with a coverslip, and observed under a fluorescence microscope (Olympus BX53 combined with U-FGW filters). Photomicrographs were captured using a 3CCD digital camera (Olympus DP74).

### (h) Assessing sperm viability and motility after preservation under aerobic and anoxic conditions

To investigate the positive effect of anoxia on sperm viability, we exposed sperm cells to 40 μL of 39.44 mM sodium sulphite in PBS (anoxic condition) and PBS (aerobic condition) for 6 h, 24 h, 7 d, and 10 d at 23 °C. A 0 h control was also prepared by staining sperm samples immediately after introducing them to PBS. Because 39.44 mM sodium sulphite cannot maintain anoxia in a solution for more than 24 h, sperm samples preserved in anoxic conditions for 24 h, 7 d, and 10 d were kept in a plastic bag with an AnaeroPack kenki pouch soon after sperm cells were prepared in anoxic PBS created by adding sodium sulphite. Further, the live and dead sperm cells were stained green and red fluorescence respectively using LIVE/DEAD^™^ Sperm Viability Kit (Thermo Fisher Scientific). Ten microlitre SYBR-14 working solution diluted 50-fold with each solution was added into 10 μL out of the 40 μL sperm suspension and incubated for 10 min, and 4 μL propidium iodide was added and incubated for 7 min under dark condition, following the methods of den Boer et al. [9]. Three microlitre of the solution was placed on a slide and covered with a coverslip. Sperm cells preserved for 0 h, 6 h and 24 h were observed using a differential interference contrast microscope to confirm the position of sperm, and a fluorescence microscope (Olympus BX53 combined with U-FBNA, U-FBW, and U-FGW filters) was used to detect the live and dead cells. Photomicrographs were captured using a 3CCD digital camera (Olympus DP74). Fluorescence from sperm cells preserved for 7 d and 10 d was observed and captured using an EVOS^™^ M7000 Imaging system with GFP and RFP light cubes (Thermo Fisher Scientific). The observation was repeated four times per sperm sample, and 150–1000 sperm cells were captured within 30 min of staining with propidium iodide. Because many sperm cells were degraded in the 7 d and 10 d aerobic condition exposures, no sperm cells were observed in two and three of the six sperm samples, and only 15–477 sperm cells were observed in the remaining sperm samples. The number of living and dead sperm cells was counted on a computer monitor. Because there were few dual-stained sperm cells, they were excluded from the assessment of sperm viability.

We also confirmed whether sperm cells preserved under anoxic PBS conditions using sodium sulphite and AnaeroPack kenki pouch (for 1, 2, 4, 8, 18, and 24 h) could be motile after exposure to aerobic conditions. A 0.5-mm deep silicone rubber spacer was placed on the slide glass to keep space between the slide and coverslip. Five microlitres of the sperm sample in anoxic sodium sulphite solution was deposited on a slide glass and covered with a coverslip. After confirmation of their immobility, 30 μL of aerobic PBS was added, and sperm motility was observed under a differential interference contrast microscope (Olympus BX53), photomicrographed using a 3CCD digital camera (Olympus DP74).

### (i) Statistical analysis

All statistical analyses were performed using R version 4. 1. 2 [10].

## 3. Results

### (a) Oxygen concentrations in the spermathecal reservoir and other tissues

Oxygen concentrations in the spermathecal reservoir of *C. osakensis* and *L. hayashi* queens were 0.284 ± 0.036% (mean ± SD) and zero, respectively (figure 1*a* and 1*b*). This was extremely low compared with those of other tissues. Oxygen concentrations not only in the spermathecal reservoir (mean ± SD = 0.211 ± 0.189%) but in the bursa copulatrix, thorax, and abdomen were extremely low in *C. osakensis* queens soon after the nuptial flight (figure 1*c*).

**Figure 1.**
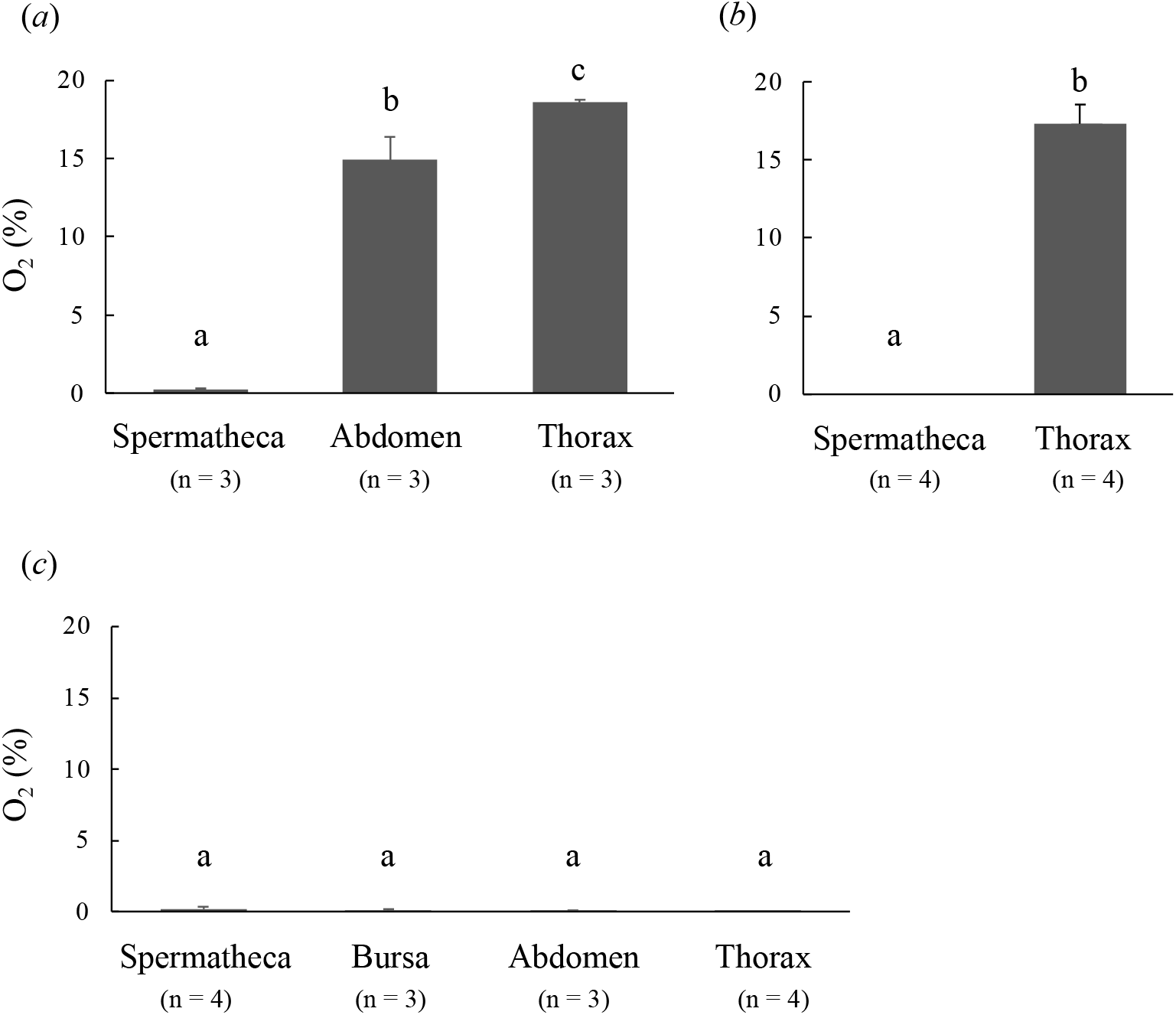
Tissue oxygen concentration in (*a*) *Crematogaster osakensis* queens at 1 year, (*b*) *Lasius hayashi* queens at 7 months after the beginning of sperm storage (or nuptial flight), and (*c*) *C. osakensis* queens soon after nuptial flight. Error bars represent standard deviation. Different alphabetical symbols indicate statistical significance at *p* < 0.05 based on Tukey’s HSD in (*a*) and (*c*) and Mann– Whitney *U* test in (*b*).

### (b) Artificial anoxic conditions induce sperm immobilisation

Sperm cells began swimming immediately after they were exposed to the aerobic PBS control (n = 5), whereas they were immotile soon after they were exposed to the hypoxic PBS and began swimming a 2–4 min later (n = 4, movie 1).

Unlike in the aerobic PBS control, complete sperm immobilisation was observed when sodium sulphite was used for creating anoxic conditions (figure 2 and movie 2). However, the sperm cells began swimming when, 5 min later, double the quantity of aerobic PBS buffer was added to the immotile sperm cells. This indicated that the sperm cells were not dead, but only immobilised under anoxic conditions. Furthermore, in both the oxygen-saturated sodium sulphite and PBS solutions, sperm cells were motile, indicating that adding sodium sulphite to PBS does not affect sperm immobilisation (figure 2).

**Figure 2.**
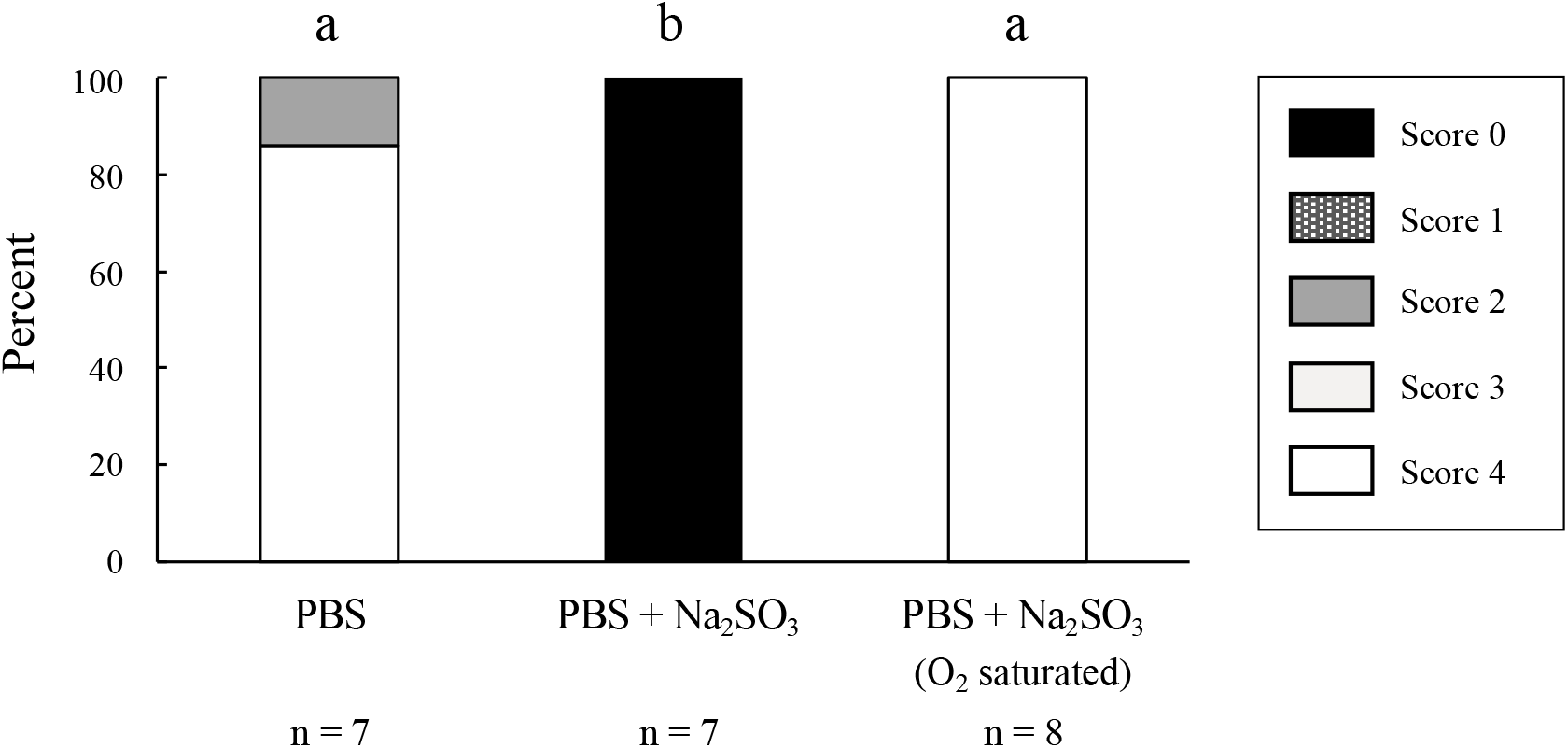
Sperm motility in the aerobic PBS control, the anoxic PBS condition created by adding sodium sulphite and an oxygen-saturated sodium sulphite solution to PBS. Different alphabetical symbols indicate statistical significance at *p* < 0.05, based on Tukey’s HSD.

### (c) Effect of glycolytic and mitochondrial respiratory pathway inhibitors on sperm motility

Sperm motility was suppressed by mitochondrial respiratory inhibitors—antimycin, oligomycin, and FCCP—although only FCCP had a statistically significant effect. The glycolytic pathway inhibitors heptelidic acid and iodoacetic acid did not affect sperm motility (figure 3).

**Figure 3.**
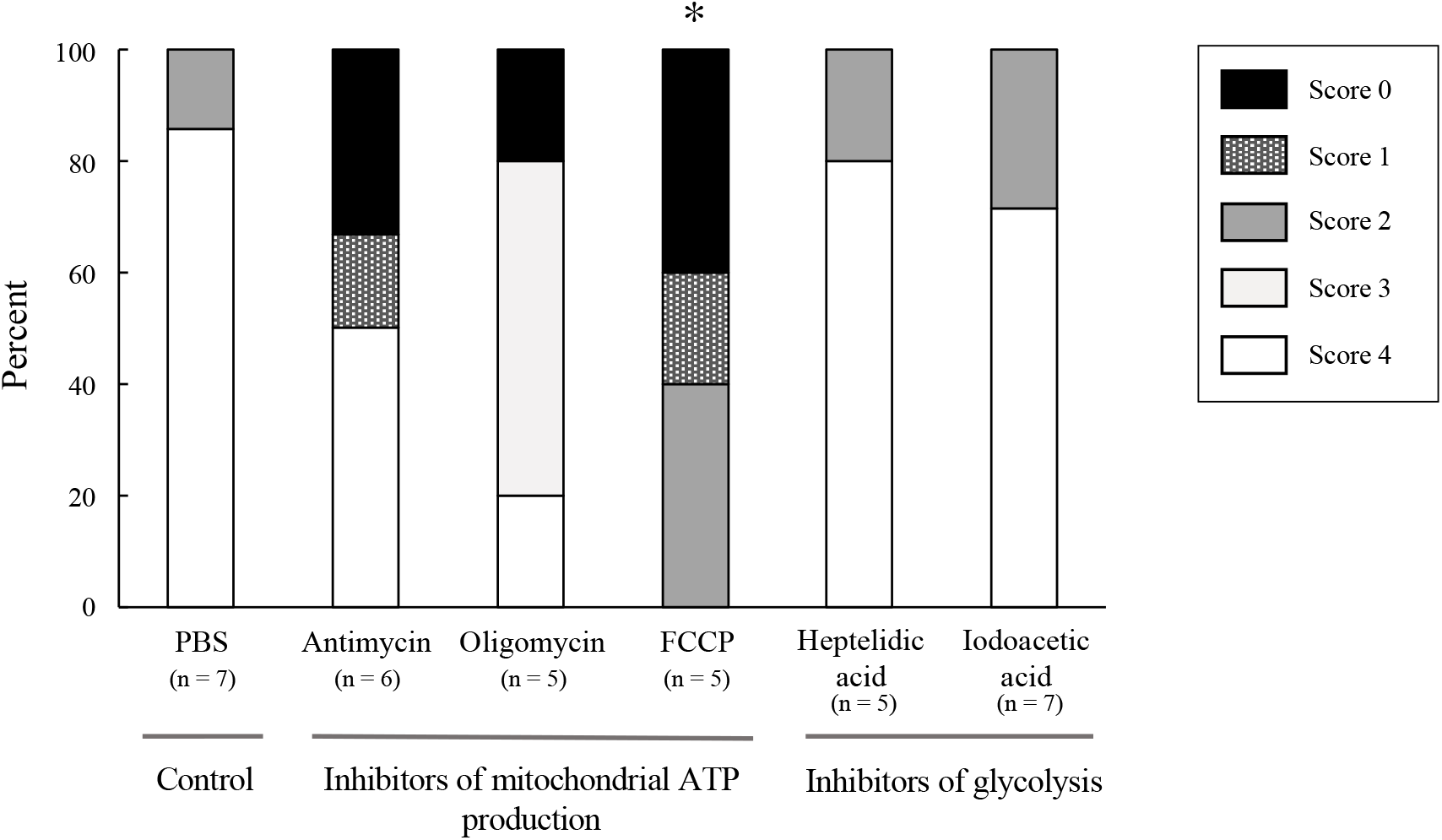
Effect of mitochondrial ATP production and glycolysis inhibitors on sperm motility. The asterisk indicates a statistically significant difference compared with the control at *p* < 0.05, based on a Kruskal–Wallis *H* test and Steel’s multiple comparison Wilcoxon tests.

### (d) Sperm cellular status under anoxic and aerobic conditions

Motile sperm cells induced by PBS did not produce more ATP than immobilised sperm cells exposed to sodium sulphite (figure 4), and membrane potentials of sperm mitochondria were similar under the anoxic and aerobic conditions (figure 5).

**Figure 4.**
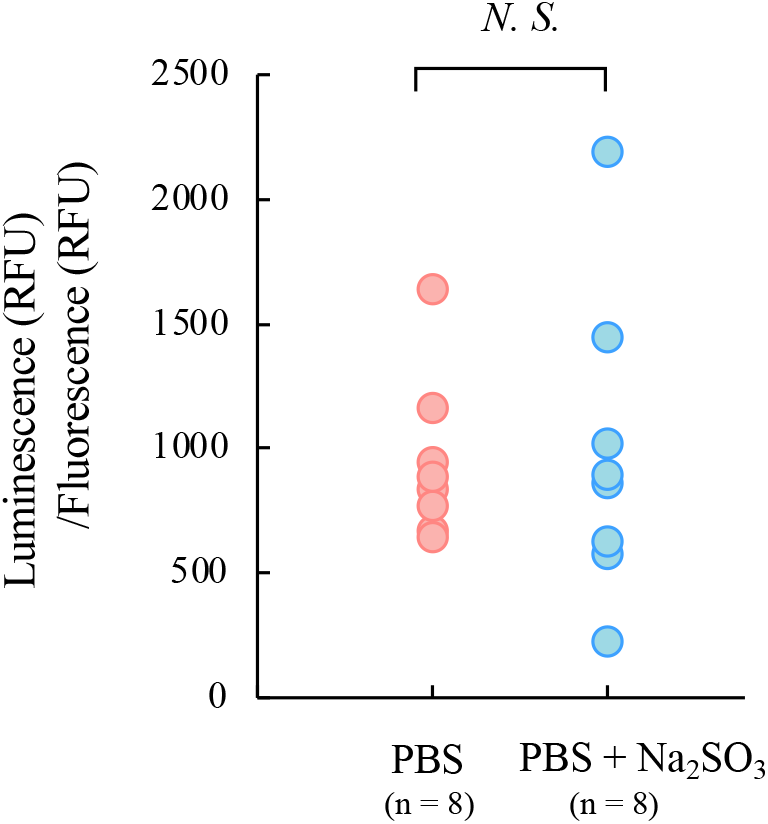
Sperm ATP levels calibrated by the number of cells in the aerobic PBS control and anoxic PBS (created by adding sodium sulphite).

**Figure 5.**
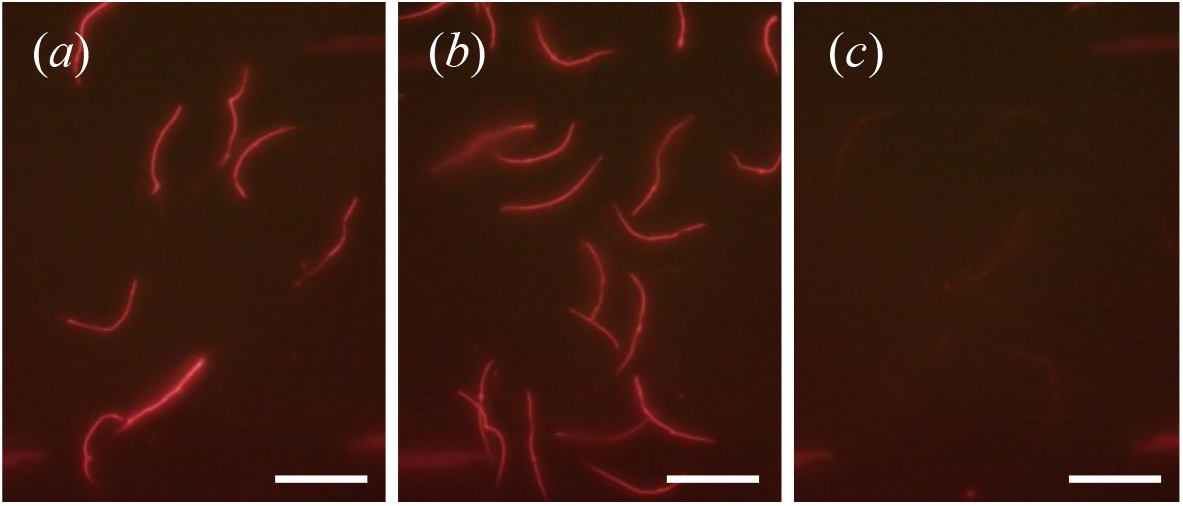
Mitochondrial membrane potential of sperm in (*a*) aerobic PBS control, (*b*) anoxic PBS (after adding sodium sulphite), and (*c*) aerobic PBS containing buffer FCCP as negative control. Scale bars indicate 50 μm.

### (e) The pH of spermathecal fluid and its effect on sperm motility

The pH of spermathecal fluid was determined to assess whether it presents an acidic environment under anoxic conditions. The pH in the reservoir was 8.5, which was higher than that of the haemolymph (approximately 7.0; figure 6*a*). The intensity of the fluorescence from the sperm increased in proportion to extracellular fluid pH. Furthermore, the intensity was similar in response to extracellular fluid with and without valinomycin and nigericin (which equilibrates intra- and extra-cellular pH). Therefore, extracellular pH equalised intracellular sperm pH, and the intracellular pH of the sperm preserved in the spermathecal reservoir fluid was not likely to be acidic.

**Figure 6.**
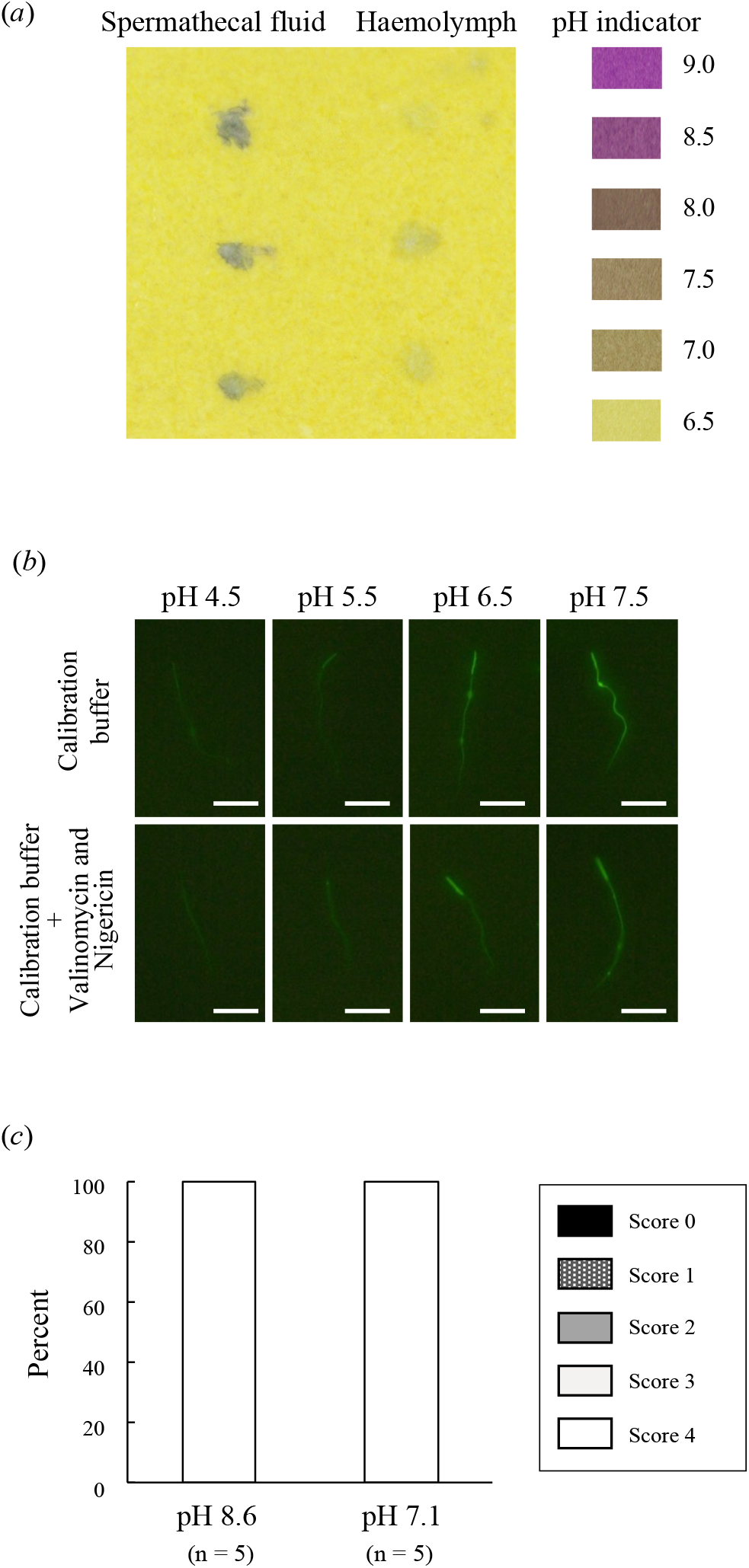
(*a*) pH of the spermathecal fluid and haemolymph. (*b*) Effect of extracellular fluid pH on intracellular pH of sperm. Fluorescence intensity of BCECF-loaded sperm exposed to various pH-calibrated buffers and those containing valinomycin and nigericin (to equilibrate the pH inside and outside cells). Scale bars indicate 20 μm. (*c*) Effect of pH-mimicked spermathecal fluid (pH 8.6) and haemolymph (pH 7.1) on sperm motility.

Sodium chloride solutions adjusted to pH 8.6 and 7.1 mimicking spermathecal reservoir fluid and haemolymph pH, respectively, did not induce sperm immobilisation (figure 6*c*).

### (f) Effect of preservation under aerobic and anoxic environments on sperm viability and physiology

The viability of sperm preserved for 6 h, 24 h, 7 d, and 10 d under anoxic conditions was higher than that under aerobic conditions (figure 7*a*). In contrast to anoxic conditions, no sperm survived, and sperm aggregation and abnormalities were observed under aerobic conditions in the 7 d and 10 d treatments (figure 7*b, c*)

**Figure 7.**
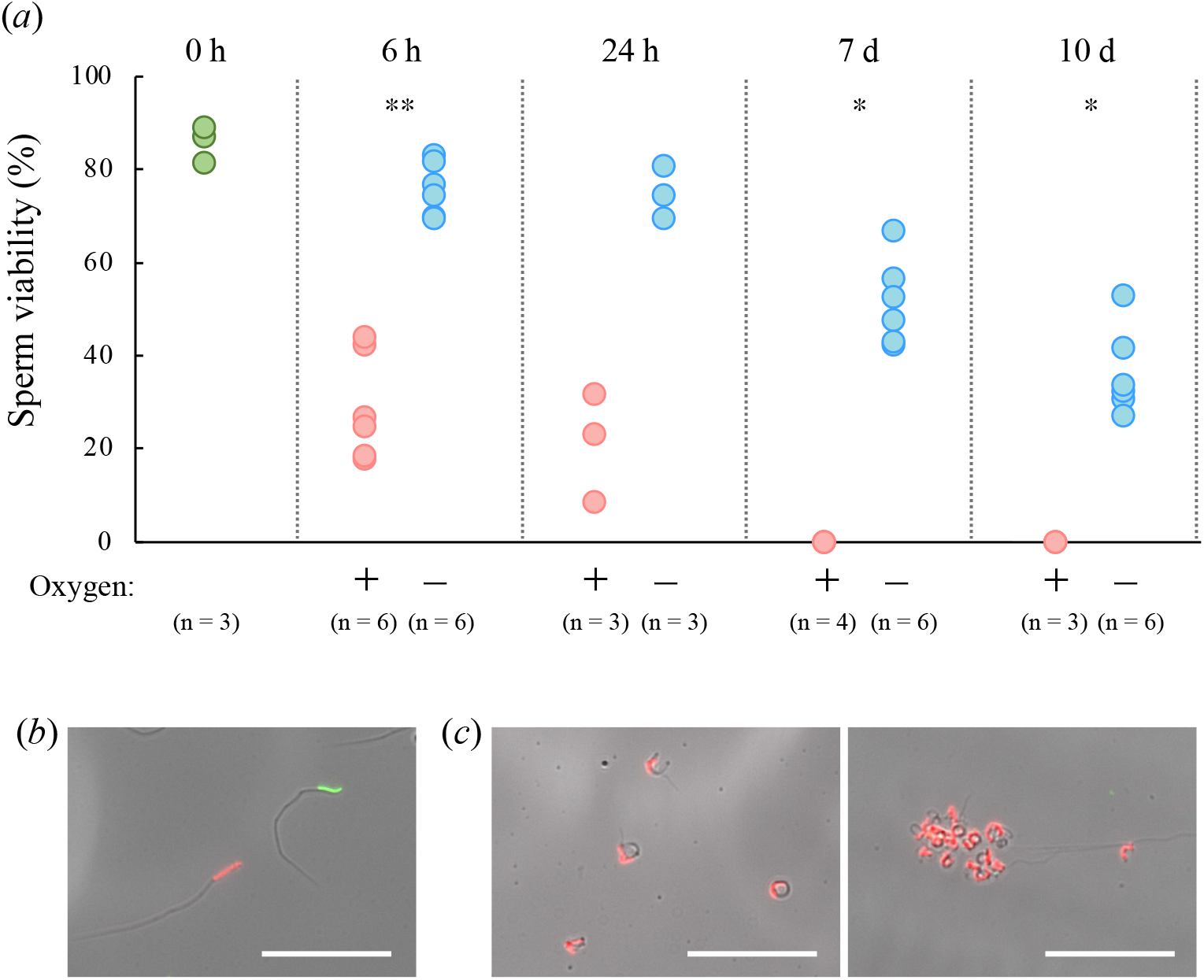
(*a*) Sperm viability under aerobic (+) and anoxic (after adding sodium sulphite) (-) PBS conditions. Asterisks indicate statistical significance based on Mann–Whitney *U* tests (* *p* < 0.05, \*\**p* < 0.005). (*b*) Image of surviving sperm stained green and dead sperm stained red under artificially anoxic conditions for 7 d. (*c*) Images of dead sperm cells with abnormal shape (left), and sperm aggregation (right). Scale bars indicate 50 μm.

Sperm cells preserved under artificial anoxic conditions for 1 and 2 h began swimming when aerobic PBS was added. However, we observed no motility in the 8, 18, and 24 h preservations, and very few were motile in the 4 h preservation (n = 3, movie 3).

### 4. Discussion

### (a) Sperm cells are stored in near-anoxic conditions and remain immotile

In this study, we found that the oxygen concentration in the spermathecal reservoir was extremely low in the two ant species, particularly in *L. hayashi* queens. Inserting oxygen probes into the small spermathecal reservoirs of *C. osakensis* queens was technically difficult, and oxygen could diffuse into the spermathecal fluid during the insertion. Therefore, the true oxygen concentration may be lower than our data reflects. This indicates that they have a powerful oxygen removal system inside the spermathecal reservoir. Because the two species belong to different subfamilies, the near-anoxic conditions in the spermathecal reservoir may be a general feature in ants.

We also demonstrated, using two methods, that artificial hypoxic and anoxic environments induced sperm immobilisation. Although many studies have examined sperm motility in artificial fluids in insects, it has been challenging to confirm that sperm motility actually occurs *in vivo* because of the difficulty in measuring the environment around sperm in the small reproductive tract [11]. Therefore, this study is one of the few that link the internal environment of the reproductive organ and the effect of the factor on sperm motility by using ant queens with a large spermatheca and a pronounced sperm storage ability.

### (b) Sperm immobilization mechanism during sperm transfer into the spermathecal reservoir

Our previous study revealed that sperm cells are immotile when they are transferred to the spermathecal reservoir of the queen soon after copulation [5]. In this study, we have demonstrated that the spermathecal reservoir and bursa copulatrix and the non-reproductive tissues, such as the abdomen and thorax, in ant queens were nearly anoxic soon after the nuptial flight. Consequently, sperm immobilisation during its transfer is presumed to be caused by the near-anoxic conditions throughout the body. Ant queens probably use their flight muscles intensively during the nuptial flight. Thus, the anoxic conditions throughout the body may result from oxygen consumption during that flight. However, there are also ant species in which the queen copulates with males around their natal nest without nuptial flight [12]. Thus, it is also necessary to assess the oxygen concentration and sperm motility soon after their copulation in these species.

It is unclear whether sperm cells are motile after they are released from the spermathecal reservoir for fertilisation [5]. It was technically challenging to measure oxygen concentration inside the spermathecal duct and bursa copulatrix because of their narrow lumens; therefore, we could not assess sperm motility for fertilisation in terms of the oxygen level of the immediate environment.

### (c) Mechanism of sperm immobilisation under anoxic conditions

It was considered that ATP for sperm motility was produced by oxidative phosphorylation through aerobic respiration in animal species; however, species such as the insects *Tribolium castaneum* [13], *Aleochara bilineata* [14] and the mammalian mouse [15] have been found to use a glycolytic system. In this study, sperm motility was restricted by respiratory inhibitors, not by glycolytic pathway inhibitors. Aerobic respiration through glycolysis, the tricarboxylic acid cycle, and oxidative phosphorylation together yields 38 mol of ATP, whereas glycolysis alone generates only 2 mol of ATP per mol of glucose. Sperm immobilisation in the spermathecal reservoir of an ant queen may be caused by a shortage of ATP generated from only glycolysis. However, we did not find more intracellular ATP under aerobic than that under anoxic conditions. It may be caused by balance between ATP consumption for sperm motility and production under aerobic conditions. A mitochondrial membrane potential was detected in the sperm under anoxic conditions. It has various roles besides ATP production, such as antiviral signalling [16] and generating a pulling force for translocating newly synthesised proteins into the mitochondrial matrix [17]. In ant sperm, elongated mitochondria occupy a large part of the flagellum as well as in other insects [11, 18] although ant sperm cells are stored in nearly-anoxic conditions. Therefore, the presence of a mitochondrial membrane potential in nearly-anoxic conditions is necessary for sperm functions, which requires further research.

Usually, either exogenous or endogenous substrates, such as sugars and lipids, are necessary as an energy resource for sperm motility. The ant sperm cells ejected from spermathecae were motile in the PBS buffer, which did not contain sugars and lipids, indicating the presence of an intracellular endogenous substrates. Also, the glycolytic pathway inhibitors heptelidic acid and iodoacetic acid, which inhibit catalysis of glyceraldehyde 3-phosphate to 1,3-bisphosphoglyceric acid by glyceraldehyde 3-phosphate dehydrogenase, did not induce sperm immobilisation. This suggests that sperm cells accumulate energy substrates in the pathway after the point of 1,3-bisphosphoglyceric acid in glycolysis and the tricarboxylic acid cycle.

### (d) Sperm immobilisation is not related to pH of the spermathecal fluid

Female Japanese quails can store sperm for 11–12 days in sperm storage tubules after mating, and stored sperm cells are also immobilised in the hypoxic environment of the sperm storage tubules [8]. However, the sperm immobilisation process is different from ants. In Japanese quails, lactic acid is accumulated through the glycolytic pathway under the hypoxic conditions of the sperm storage tubules, which decreases the intracellular sperm pH. Because the dynein ATPase activity of sperm is generally inhibited by acidic pH, the sperm of Japanese quails cannot swim while in the sperm storage tubules [8, 11]. However, our study revealed that the pH of the spermathecal fluid was high (approximately 8.5) and that intracellular sperm pH was related to extracellular pH in ants. Therefore, sperm immobilisation could not be caused by sperm acidification. This also suggests that the spermathecal reservoir regulates luminal pH and eventually inhibits acidification of stored sperm in a nearly anoxic environment. In ant queens, the columnar epithelial cells of the spermathecal reservoir are considered to have transport functions [3], and in cancer cells transporters play an important role in inhibiting lactic acid accumulation under hypoxic conditions [19]. Therefore, a pH regulation system mediated by transporters and channels related to transition protons and lactic acid should exist in both the spermathecal epithelium and sperm plasma membrane.

High intracellular pH, which mimicked the spermathecal fluid pH, did not induce sperm immobilisation, indicating that a high pH does not affect sperm motility in ants. In this experiment, a NaCl solution, which has simpler contents than PBS, could induce sperm motility, suggesting that these ion components are also not related to the induction of sperm motility.

### (e) Evolution of a sperm storage system in Hymenoptera

The order Hymenoptera is a large group, and sperm storage duration ranges from several weeks or months in solitary species to several decades in ants that have developed prominent eusociality. Honeybee (*Apis mellifera*) queens can store sperm cells for 2–4 years [20] which is a long duration, second only to ants— although they belong to a different lineage. Near anoxia, high pH of the spermathecal fluid, and immobility of stored sperm in *A. mellifera* queens have been reported [21-23]. It is suggested that the two lineages evolved a common mechanism for sperm maintenance. In honeybee queens, the spermathecal reservoir is surrounded by a tracheal network [24-26], which is considered to suppress oxygen spread to the spermathecal reservoir and maintain near anoxia [23]. However, ant spermathecae possess no tracheal network [3]. In both honeybees and ants, abundant mitochondria are present in the columnar epithelium of the spermathecal reservoir [3]; therefore, they may consume oxygen and maintain near anoxia inside the reservoir. To our knowledge, studies investigating sperm physiology and molecular mechanisms of spermathecae are limited except for those on honeybees and ants, which show long-term sperm storage by queens. Future studies should compare sperm maintenance systems among solitary and social hymenopteran species with short to long sperm storage durations, and conduct phylogenetic analyses to understand the evolution of sperm storage ability in Hymenoptera.

### (f) Anoxic environment improves sperm viability

In this study, we clearly showed that a near-anoxic environment is crucial for sperm viability. The anoxic conditions and consequent sperm immobilisation should have significant advantages for sperm physiology in reducing risks from the production of reactive oxygen species and physical cell damage. It is, however, still unknown whether sperm viability was enhanced by sperm immobilisation alone or through pleiotropic effects of the anoxic condition. This should become clear when the requirements for sperm immobilisation under aerobic conditions are discovered.

Under hypoxic conditions, reactive oxygen species are generated by excess electrons because oxygen is the ultimate electron acceptor in the mitochondrial electron transport system [27]. In ant and honeybee spermathecae, the genes and proteins related to antioxidant function are enriched [28-31]. The antioxidant system may, therefore, protect against oxygen occurring accidentally in the spermathecal reservoir.

Interestingly, sperm cells preserved in an artificial anoxic solution induced by sodium sulphite for more than 4 h differed from those preserved for 2 h because the 4 h-anoxic, but not the 1 and 2 h-anoxic, most of sperm cells did not begin swimming when aerobic PBS was introduced. This indicates that the physiology of sperm motility was altered within 4 h under anoxia, yet the survival rate was still high. Furthermore, the longer sperm cells were preserved under anoxia, the lower their viability gradually became, where the viability of sperm cells preserved for 0 h and 10 d was > 80% and ∼ 30%, respectively. This indicates that the near anoxic condition is a key factor for long-term sperm survival, but other factors may also be essential. Therefore, for a complete understanding of long-term sperm storage mechanisms in ant queens, future studies on the role of other factors—such as ions, metabolites, and proteins [31] —are necessary.

## Supplementary information

**movie 1**

Sperm motility under hypoxic PBS conditions (created using AnaeroPack^®^ kenki pouch) and aerobic PBS control.

**movie 2**

Sperm motility in aerobic PBS control and anoxic PBS conditions (created by adding sodium sulphite).

**movie 3**

Motility of sperm cells preserved in anoxic conditions for 2, 4, and 8 h and subsequently exposed to aerobic PBS solutions.

## Authors’ contributions

All authors designed the studies. AG and MT collected and analyzed the data. AG wrote the manuscript. All authors have read and approved the final version of the manuscript.

## Acknowledgments

We thank members of laboratory of entomology in Kagawa University for their assistance with the sample collection. We are also grateful to Dr. T. Sasanami, Dr. M. Matsuzaki and Dr. K. Takeda for useful comments.

## References

1. Orr, T.J. & Zuk, M. 2012 Sperm storage. Curr. Biol. 22, R8–R10. (doi:10.1016/j.cub.2011.11.003).

2. Keller, L. 1998 Queen lifespan and colony characteristics in ants and termites. Ins. Soc. 45, 235–246. (doi:10.1007/s000400050084).

3. Wheeler, D.E. & Krutzsch, P.H. 1994 Ultrastructure of the spermatheca and its associated gland in the ant Crematogaster opuntiae (Hymenoptera, Formicidae). Zoomorphology 114, 203–212. (doi:10.1007/BF00416859).

4. Gobin, B., Ito, F., Peeters, C. & Billen, J. 2006 Queen-worker differences in spermatheca reservoir of phylogenetically basal ants. Cell. Tissue Res. 326, 169–178. (doi:10.1007/s00441-006-0232-2).

5. Gotoh, A. & Furukawa, K. 2018 Journey of sperms from production by males to storage by queens in Crematogaster osakensis (Hymenoptera: Formicidae). J. Insect Physiol. 105, 95–101. (doi:10.1016/j.jinsphys.2018.01.008).

6. Osanai, M., Kasuga, H. & Aigaki, T. 1989 Induction of motility of apyrene spermatozoa and dissociation of Eupyrene sperm bundles of the silkmoth, Bombyx mori, by initiatorin and trypsin. Inver. Reprod. Dev. 15, 97–103. (doi:10.1080/07924259.1989.9672029).

7. Alavi, S.M.H. & Cosson, J. 2006 Sperm motility in fishes. (II) Effects of ions and osmolality: A review. Cell Biol. Int. 30, 1–14. (doi:10.1016/j.cellbi.2005.06.004).

8. Matsuzaki, M., Mizushima, S., Hiyama, G., Hirohashi, N., Shiba, K., Inaba, K., Suzuki, T., Dohra, H., Ohnishi, T., Sato, Y., et al. 2015 Lactic acid is a sperm motility inactivation factor in the sperm storage tubules. Sci. Rep. 5, 17643. (doi:10.1038/srep17643).

9. den Boer, S.P.A., Boomsma, J.J. & Baer, B. 2008 Seminal fluid enhances sperm viability in the leafcutter ant Atta colombica. Behav. Ecol. Sociobiol. 62, 1843–1849. (doi:10.1007/s00265-008-0613-5).

10. Team, R.C. 2021 R: A Language and Environment for Statistical Computing. R Foundation for Statistical Computing, Vienna, Austria.

11. Werner, M. & Simmons, L.W. 2008 Insect sperm motility. Biol. Rev. Camb. Philos. Soc. 83, 191–208. (doi:10.1111/j.1469-185X.2008.00039.x).

12. Hölldobler, B. & Wilson, E.O. 1990 The ants. Cambridge, Mass., Harvard University Press.

13. Bloch Qazi, M.C., Aprille, J.R. & Lewis, S.M. 1998 Female role in sperm storage in the red flour beetle, Tribolium castaneum. Comp. Biochem. Physiol. Part A Mol. Integr. Physiol. 120, 641–647. (doi:10.1016/S1095-6433(98)10081-8).

14. Werner, M., Zissler, D. & Peschke, K. 1999 Structure and energy pathways of spermatozoa of the rove beetle Aleochara bilineata (Coleoptera, Staphylinidae). Tissue Cell 31, 413–420. (doi:10.1054/tice.1999.0052).

15. Mukai, C. & Okuno, M. 2004 Glycolysis plays a major role for adenosine triphosphate supplementation in mouse sperm flagellar movement. Biol. Reprod. 71, 540–547. (doi:10.1095/biolreprod.103.026054).

16. Koshiba, T., Yasukawa, K., Yanagi, Y. & Kawabata, S. 2011 Mitochondrial membrane potential is required for MAVS-mediated antiviral signaling. Sci. Signal 4, ra7. (doi:10.1126/scisignal.2001147).

17. Sato, T.K., Kawano, S. & Endo, T. 2019 Role of the membrane potential in mitochondrial protein unfolding and import. Sci. Rep. 9, 7637. (doi:10.1038/s41598-019-44152-z).

18. Wheeler, D.E., Crichton, E.G. & Krutzsch, P.H. 1990 Comparative ultrastructure of ant spermatozoa (formicidae: Hymenoptera). J. Morphol. 206, 343–350. (doi:10.1002/jmor.1052060311).

19. Rademakers, S.E., Lok, J., van der Kogel, A.J., Bussink, J. & Kaanders, J.H.A.M. 2011 Metabolic markers in relation to hypoxia; staining patterns and colocalization of pimonidazole, HIF-1α, CAIX, LDH-5, GLUT-1, MCT1 and MCT4. BMC Cancer 11, 167. (doi:10.1186/1471-2407-11-167).

20. Dade, H.A. 1994 Anatomy and Dissection of the Honeybee, International Bee Research Association.

21. Lensky, Y. & Schindler, H. 1967 Motility and reversible inactivation of honeybee spermatozoa in vivo and in vitro. Ann. Abeille. 10, 5–16.

22. Gessner, B. & Gessner, K. 1976 Inorganic ions in spermathecal fluid and their transport across the spermathecal membrane of the queen bee, Apis mellifera. J. Insect Physiol. 22, 1469–1474. (doi:10.1016/0022-1910(76)90212-2).

23. Paynter, E., Millar, A.H., Welch, M., Baer-Imhoof, B., Cao, D. & Baer, B. 2017 Insights into the molecular basis of long-term storage and survival of sperm in the honeybee (Apis mellifera). Sci. Rep. 7, 40236. (doi:10.1038/srep40236).

24. Poole, H.K. 1970 The wall structure of the honey bees spermatheca with comments about Its function. Ann. Entomol. Soc. Am. 63, 1625–1628. (doi:10.1093/aesa/63.6.1625).

25. Dallai, R. 1975 Fine structure of the spermatheca of Apis mellifera. J. Insect Physiol. 21, 89–109. (doi:10.1016/0022-1910(75)90072-4).

26. Gotoh, A. & Sasaki, K. 2021 Caste differentiation of spermatheca and organs related to sperm use and oviposition in the honeybee, Apis mellifera. Apidologie 52, 262–271. (doi:10.1007/s13592-020-00815-9).

27. Fukuda, R., Zhang, H., Kim, J.W., Shimoda, L., Dang, C.V. & Semenza, G.L. 2007 HIF-1 regulates cytochrome oxidase subunits to optimize efficiency of respiration in hypoxic cells. Cell 129, 111–122. (doi:10.1016/j.cell.2007.01.047).

28. Weirich, G.F., Collins, A.M. & Williams, V.P. 2002 Antioxidant enzymes in the honey bee, Apis mellifera. Apidologie 33, 3–14.

29. Collins, A.M., Williams, V. & Evans, J.D. 2004 Sperm storage and antioxidative enzyme expression in the honey bee, Apis mellifera. Insect Mol. Biol. 13, 141–146. (doi:10.1111/j.0962-1075.2004.00469.x).

30. Baer, B., Eubel, H., Taylor, N.L., O’Toole, N. & Millar, A.H. 2009 Insights into female sperm storage from the spermathecal fluid proteome of the honeybee Apis mellifera. Genome Biol. 10, R67. (doi:10.1186/gb-2009-10-6-r67).

31. Gotoh, A., Shigenobu, S., Yamaguchi, K., Kobayashi, S., Ito, F. & Tsuji, K. 2017 Transcriptome profiling of the spermatheca identifies genes potentially involved in the long-term sperm storage of ant queens. Sci. Rep. 7, 5972. (doi:10.1038/s41598-017-05818-8).

